# The Ph.D. Panic: Examining the relationships among teaching anxiety, teaching self-efficacy, and coping in Biology graduate teaching assistants (GTAs)

**DOI:** 10.1101/2020.02.07.938597

**Authors:** Miranda M. Chen Musgrove, Elisabeth E. Schussler

**Affiliations:** Department of Ecology and Evolutionary Biology, The University of Tennessee, Knoxville, TN 37996

**Keywords:** teaching anxiety, teaching self-efficacy, coping strategies, coping frequency, Biology, graduate teaching assistants (GTAs)

## Abstract

Anxiety among graduate students in the United States has increased over the last several decades, affecting not only their overall mental health but also reducing retention in graduate programs. Teachers with high teaching anxiety can negatively impact student learning, yet the impacts of teaching anxiety on graduate teaching assistants (GTAs) is not well studied. Biology GTAs teach most introductory Biology labs and discussions nationally, thus broadly influencing the quality of undergraduate education. In Fall 2016, we investigated Biology GTA teaching anxiety at a large research-intensive southeastern university by (1) measuring teaching anxiety of Biology GTAs, and (2) exploring the relationships among teaching anxiety, self-efficacy, and coping. Using multiple linear regressions, we found that greater teaching self-efficacy is related to lower teaching anxiety in Biology GTAs (*R*^*2*^_*adi*_ =0.65, *p*<0.001). Coping strategies and frequencies did not significantly contribute to teaching anxiety in our models. We found similar levels of teaching anxiety across genders, ethnicities, student citizenship status (domestic vs. international) and teaching experience level. However, there were significant differences among student subgroups in teaching self-efficacy and coping strategies. Effective coping may contribute to the lack of anxiety differences among some of the student subgroups. These results can inform teaching professional development for GTAs, and encourage greater awareness and dialogue about the impacts of mental health issues in academia.

## INTRODUCTION

The reported incidence of anxiety in graduate students in the United States has been rising markedly over the last several decades [1]. Graduate students in the United States are six times more likely to experience depression and anxiety than the general public [2,3]. Anxiety is defined as the state of anticipatory apprehension over possible deleterious happenings [4]. Anxiety stimulates physiological responses similar to stress: increased levels of cortisol, faster heartrate, dilated pupils, etc. However, these physical changes accompany feelings of concern or worry over an anticipated event or outcome that may happen in the future [5].

Research universities depend on graduate students for instruction, especially for large enrollment classes [6]. Biology graduate teaching assistants (GTAs) have been estimated to teach over 91% of freshman Biology labs and discussions nationally [6–8]. According to the Longitudinal Study of Future STEM Scholars (LSFSS, Connolly et al. 2016), which studied more than 3,000 STEM PhD students over 4 years, nearly all (94.9%) taught undergraduates during their doctoral programs. These graduate students, however, often teach with little to no prior pedagogical training; all while establishing research projects and navigating departmental protocols and norms. As a result, GTAs may experience a lack of confidence about their teaching [10], potentially resulting in teaching anxiety. Given university reliance on GTAs for teaching, factors that decrease instructional quality (such as teaching anxiety or confidence) may greatly influence the quality of undergraduate education at the institution.

In this study, we explore factors that may impact teaching anxiety in a sample of Biology GTAs at one research institution. The literature describes theoretical relationships among anxiety, self-efficacy, and coping [4] that we test in our population, along with potential demographic and background/contextual influences. Previous studies have investigated GTAs’ self-efficacy [11,12], graduate student coping with writing [13], and GTA coping with teaching apprehension [14]. To our knowledge, this study is the first to examine Biology GTA teaching anxiety, self-efficacy, and coping under one model.

### The relationship between anxiety and self-efficacy

Bandura’s social cognitive theory, particularly pertaining to self-efficacy, provides a useful theoretical framework for studying the causes and consequences of anxiety and coping in GTAs. Social cognitive theory explains how an individual’s behavior can be shaped by personal, behavioral, and environmental influences [15]. A central concept in social cognitive theory is *self-efficacy*. Self-efficacy is the belief or confidence in one’s ability to successfully carry out a specific task or course of action [4,16]. Self-efficacy has been widely studied within psychology, and more recently within GTAs [9,11,12,17].

There are four main mechanisms that build self-efficacy: 1) mastery experiences, 2) vicarious experiences, 3) social persuasion, and 4) emotional/physiological appraisal [18]. For example, a GTA with many years of teaching experience has likely gained high teaching self-efficacy through mastery experiences. As a novice teacher, she may have chosen to observe an experienced GTA teach before running her own class—an example of building teaching self-efficacy vicariously. To improve self-efficacy through social persuasion, a GTA could be persuaded or convinced by her mentor or trusted friend that she would be a successful teacher. Lastly, the way an individual cognitively appraises an experience (positively or negatively) also influences self-efficacy for that particular situation. Powerful emotions, such as anxiety, can alter individuals’ beliefs about their capabilities. For example, a GTA nervous before entering the classroom may interpret these feelings as a sign of poor future performance, and thus have low teaching self-efficacy. All four sources of self-efficacy depend on the individuals’ cognitive processing related to the specific task, the context of said task, and self-assessment of task competence.

According to Tschannen-Moran et al. [19], *teaching self-efficacy*, specifically, is a teacher’s perception of their ability to “organize and execute courses of action required to successfully accomplish a specific teaching task in a particular context.” Teachers with stronger teaching self-efficacy beliefs often have more efficient classroom management, planning and organization, demonstrate greater enthusiasm and commitment, a greater willingness to try new pedagogical methods, persist in difficult teaching-related tasks, and is predictive of positive student achievement [20–23]. Self-efficacy is a strong predictor of success in a task; with greater self-efficacy, the more likely the task will be carried out successfully.

In the college setting, high teaching self-efficacy in GTAs correlates with strong performance in teaching [12]. Variables such as previous teaching experience, perceived quality of GTA professional development (PD), total hours of PD, and perception of the departmental climate are significant factors that impact teaching self-efficacy of STEM GTAs (DeChenne et al. 2015). Other studies suggest that participating in teaching PD significantly increases teaching self-efficacy in GTAs, particularly for women (25, 26). Therefore, we predict GTAs with high self-efficacy will have low anxiety.

### The relationship between anxiety and coping

Another factor related to anxiety is coping. Coping can be defined as an individual’s behavioral response(s) to external stressors, often with the objective to reduce or tolerate the stress [26–30]. Roach [14] described coping as “trying to find some way to deal with or address [felt] needs or problems.” Coping can be conceptualized as either (1) *adaptive* or (2) *maladaptive* [30]. Adaptive coping helps to advance individuals through problems and support their well-being (e.g. practice for a presentation, seek social support); while maladaptive coping prevents stressors or problems from being resolved and can exacerbate threats to well-being (e.g. avoid writing tasks, social withdrawal). Coping varies with the stressor, and some situations can involve both adaptive and maladaptive coping (e.g. returning to writing tasks after initial avoidance). With Biology GTAs balancing multiple roles as teachers, researchers, students, and employees, those with anxiety need effective coping strategies. We predict that higher coping and use of effective coping strategies should lead to lower anxiety. Those with higher anxiety, may not coping or not using effective coping strategies. Within the GTA literature, there has been little research examining coping efficacy and frequency in relation to anxiety.

Investigating anxiety under this framework, we recognize the implicit assumption that anxiety is a negative emotion. However, Yerkes and Dodson [31] established a threshold in which “arousal” or anxiety can actually increase productivity, and Pekrun et al. [5] acknowledged its ability to be an activating emotion in terms of motivation. Pelton [32] suggested that reducing *all* sources of anxiety is not ideal. Thus, too much or too little anxiety about teaching are both likely impediments to teaching effectiveness, and to a certain extent, some level of doubt or lack of confidence may provide the impetus to improve teaching effectiveness [33].

Though we recognize some anxiety as motivating, anxiety can only be productive if it is paired with constructive and effective coping strategies. If the coping is maladaptive and destructive, anxiety may have a negative impact. We predict that higher coping and use of effective coping strategies should lead to lower anxiety. Those with higher anxieties, may not be coping or not using effective coping strategies.

### Background influences: Demographics and Context

Anxiety is not homogenous in a population, and teaching anxiety is presumably not as well. Some groups of GTAs (e.g. domestic, experienced) may be better able to cope with teaching anxiety than other groups, thereby leading to lower reported anxieties [2,34]. Differential levels of anxiety can be the result of several factors, such as teaching experience level [35], gender [2], or student citizenship status or nationality [34]. Because of this, teaching anxiety will likely differ between some graduate student sub-populations (such as genders, racial/ethnic groups, novice vs. experienced GTAs, international vs. domestic GTAs, etc.); thus, different groups of students may require unique coping strategies and resources. For example, women and other minority groups suffer differential impacts of mental distress, with 43% and 41% of women in graduate school reporting anxiety and depression, respectively, compared to 34% and 35% of men [2]. International students in the United States also report different academic challenges compared to their domestic counterparts, such as concern over program structure, career preparation, and alignment with career goals [34]. Individuals with generalized anxiety disorder may also represent a unique population in terms of teaching anxiety. When studying anxiety in any context, it is important to capture contextual and demographic variables that may account for differences in anxiety in certain subgroups of the study population.

### Research questions

This study investigated graduate student teaching anxiety in Biology GTAs at a large research-intensive university in Fall 2016. We collected and analyzed data to answer two research questions:

1. In what ways do GTAs (and certain subgroups of GTAs) differ in teaching anxiety, teaching self-efficacy, coping strategies, and coping frequencies?
2. How do GTA teaching self-efficacy, coping, and contextual variables relate to teaching anxiety and each other?

Answering these questions will reveal how teaching anxiety may vary across a population of GTAs in one disciplinary area and inform teaching professional development to help build teaching self-efficacy and effective coping strategies among GTAs.

## METHODS

### Study Population

Biology GTAs at a large research-intensive southeastern university were the study population. The GTAs were recruited from across the Division of Biology via a listserv of graduate students from three departments (Ecology & Evolutionary Biology, Microbiology, Biochemistry & Cellular and Molecular Biology) and one program (Genome Science & Technology). Of these, 211 graduate students were enrolled in a Masters or PhD program. As of Fall 2016, approximately 94% of graduate students were seeking PhDs, and 55% identified as female.

### Data Collection

In Fall 2016, an online survey was created, approved by the Institutional Review Board (IRB-16-03235-XP), and deployed to Biology graduate students via the Qualtrics survey software. The e-mail targeted individuals who were either currently teaching or who had been a GTA previously. The survey was open for two weeks at the end of October 2016. We chose mid-semester to avoid capturing anxieties related to the beginning of the semester and give GTAs time to acclimate to their multiple responsibilities that semester. To encourage participation in the survey, a small monetary compensation of $5 was offered to each responding graduate student. All relevant data are within the manuscript and its Supporting Information files (**S1 Table**).

Three validated instruments from the literature were included in the survey to measure teaching anxiety, teaching self-efficacy, and coping [11,14,36]. There were a total of 103 questions (items) in the survey, with 29 measuring teaching anxiety [36], 18 measuring teaching self-efficacy [11], and 24 measuring frequency of the enactment of coping strategies [14] (see **S2 Table** for complete survey).

*Teaching anxiety* was measured using Parson’s 29-item survey [36], which was initially developed to measure teaching anxiety in preservice K-12 teachers. The survey was further adapted for our study population (GTAs) by changing verbiage addressing “preservice teachers” to “GTAs”. Participants rated each statement on a 1-5 Likert scale, where 1 is “Never” and 5 is “Always”. For example, one item states, “I feel secure with regard to my ability to keep a class under control.” Other items probed GTAs’ feelings about having control in the classroom, answering student questions, comparing one’s abilities to others’ teaching, etc.

*Self-efficacy* was measured using DeChenne et al.’s self-efficacy survey [11], which was developed and used with a GTA population at another institution. The survey is an 18-item instrument and items are rated on a 1-5 Likert scale, with 1 being “Not confident at all” and 5 being “Very confident” [11]. Two constructs of teaching self-efficacy were measured via this survey: *learning environment self-efficacy* and *instructional self-efficacy. Learning environment self-efficacy* (11 items) is related to a teacher’s belief in being able to promote a positive learning environment via student participation, encouraging students to ask questions, engaging students to interact with each other, etc. A teacher’s *instructional self-efficacy* (7 items) is related to their confidence in being able to carry out “instructional tasks,” e.g. a teacher’s belief in being able to evaluate and assess accurately students’ academic progress, grading appropriately, clearly identify learning objectives, preparedness to teach, etc.

*Coping* was measured using Roach’s instrument [14] that measures the frequency of six types of coping strategies in response to teaching anxiety: (1) *preparing materials*, (2) *muscular desensitization* e.g. breathing deeply or muscular exercises, (3) *cognitive restructuring* e.g. positive thinking, (4) *preparing delivery*, (5) *visualization* e.g. imagining successfully teaching the class, and (6) *mentoring*, e.g. reaching out to other GTAs or faculty. This instrument was developed for GTAs across multiple disciplines and countries of origin to measure how GTAs reduce anxiety in preparation for teaching their class. Instrument items were based on techniques for coping with communication apprehension [14]. It has 24 items with at least two items per construct. Participants rate the frequency of their coping activities on a 1-5 Likert scale, ranging from 1 “Never” to 5 “Always” before teaching. For example, an item from type 3 coping asks participants to rate how often they “practice saying and thinking positive self-thoughts about yourself.”

#### Contextual variables

Lastly, there were 32 investigator-created questions, which capture demographic and other contextual variables. Four items were to measure general anxiety (GA) among GTAs and asked participants to rate their anxiety: “*About being a graduate student/the graduate student experience*,” “*Being a TA in your most recent teaching assignment*,” “*Being a GTA generally*,” and “*In your daily life generally*.” They responded using a 1-5 Likert scale, with 1 being not anxious and 5 being very anxious. Another three items asked participants about their perceptions of teaching support from their advisor, department, and institution on a scale of 1-5, 1 being no support and 5 being very supportive of teaching. We asked participants to report the average number of hours they took to prepare for teaching each week, the number of semesters of GTA experience (>1 year of GTA experience was considered “Experienced”), and career aspirations (see **S2 Table**). Demographic variables such as gender, ethnicity, department, student citizenship status or nationality, and degree sought were also included.

### Data analysis

We calculated measures of reliability and validity to determine whether the anxiety, self-efficacy, and coping instruments accurately measured the identified variables for the GTA population. Reliability measures consistency when a testing procedure is repeated [37]; while validity is a measure of its accuracy in drawing correct inferences from survey scores [38]. Two forms of evidence were used to assure reliability and validity of the three surveys. First, each instrument was vetted for this project based on reported reliability scores from the literature. The teaching anxiety scale had a reported alpha coefficient 0.93, the self-efficacy measures an alpha score of 0.90, and the coping constructs of 0.94 [11,14,36]. We also calculated Cronbach alpha scores for our GTA population. Second, content validity of the questions were checked based on professional judgment by experts (one psychology faculty and 3 biology faculty) as to the appropriateness of the instrument for the Biology GTA population [38]. Though confirmatory factor analysis (CFA) is commonly used to validate the use of an instrument with a new population, it requires a much larger data set than we had available for this project, so we were not able to conduct this analysis [37,39].

To prepare the teaching anxiety, teaching self-efficacy, and coping item results for analysis, we followed the suggested protocol for each instrument. We summed each individual’s responses to the 29-items to result in a teaching anxiety score, with half of the items being reverse scored to adjust for positive phrasing [36]. An individual could score between 29 (low anxiety) to 145 (high anxiety) on this anxiety scale. The scores for the two self-efficacy constructs were compiled separately and averaged, allowing each participant to have two teaching self-efficacy scores (learning environment self-efficacy and instructional self-efficacy). Final scores of each self-efficacy construct ranged from 1 (low self-efficacy) to 5 (high self-efficacy) [11]. Lastly, for coping, final summed scores for each type of coping range from as low as 2 to as high as 45, depending on the type [14].

Contextual variables were processed independently from one another depending on the items. Some demographic variables were dummy coded, such as gender (1 = male, 2 = female), ethnicity (0 = non-white, 1 = white), student citizenship status (domestic = 1, international = 2), degree program (1 = MS, 2=PhD), department (1 = BCMB, 2 = EEB, 3 = GST, 4 = Micro, 5 = Other), and teaching experience (0 = Novice GTA with < 1 year of experience, 1 = Experienced GTA with 1 year or more of experience). The term ‘international student’ is defined as individuals enrolled in higher education institutions who are on temporary student visas and are often non-native English speakers. The terms, ‘domestic,’ ‘local,’ or ‘resident students’ refers to students who are native English speakers residing in their own country [40]. The investigator-created items for general anxiety and perceptions of teaching support were all kept as independent items and not summed or averaged, as they were not from a validated instrument.

To address the first research question (examining the differences in anxiety among GTAs) t-tests or one-way Analysis of Variance (ANOVAs) were used to calculate differences in anxiety, self-efficacy, coping, or other survey items between subgroups (gender, ethnicity, department, citizenship, degree, experience level) of GTAs from the collected demographic data. Researcher-created general anxiety items were examined as separate items. Descriptive statistics were also calculated for each of the instruments and/or items for the entire GTA sample.

To answer the second research question (examining how anxiety, self-efficacy and coping relate) two types of statistical models were developed: bivariate correlations and multiple linear regressions (MLRs). We computed correlational analyses to examine the strength and direction of the relationships between each construct in the study. These correlations allowed us to initially explore the relationships between teaching anxiety and other constructs (self-efficacy, coping, general anxiety) and the contextual variables. Building on these analyses, we next developed multiple linear regressions (MLRs) that included self-efficacy and coping as predictors within the same model. The correlations of continuous variables and those suggested by the literature were used to inform model development. According to Tabachnick and Fidell [41], the primary goal of regression analysis is often to investigate the relationship between a dependent variable and several independent variables. Here, we sought to identify the combined variance in teaching anxiety that was accounted for when considering multiple independent variables (e.g. teaching self-efficacy, coping, demographic, and contextual variables). The variables that were included in the initial model before step-wise selection, were the results from the 3 instruments (teaching anxiety, teaching self-efficacy, and coping), general anxiety, teaching support, hours to prepare for teaching, and demographics. All values from the instruments were z-scored for comparison. A step-wise selection procedure was used to select the variables that explained significant variance (R^2^adj).

To compare multiple models and determine the most parsimonious, a measure called Akaike Information Criterion (AIC) was calculated (Tabachnick and Fidell 2007). The AIC captures both estimated residual variance and model complexity in one statistic. If the amount of residual variance decreases, so does the AIC score. If excessive parameters are added to the model, the AIC score increases. The score must be read in comparison to other models, and the model with the lowest AIC score is considered the model that explains the greatest variance of the dependent variable, while maintaining parsimony. Within each model, variance inflation factors (VIF) were also calculated. VIF quantifies how much the variance within a model is inflated by multicollinear variables [41]. If the VIF exceeds 4, further investigation is needed. If the VIF is greater than 10, there is multicollinearity between variables that needs to be corrected [42]. All survey analyses were conducted in R [43].

## RESULTS

Eighty-nine graduate students completed the Fall 2016 survey. A total of 115 individuals attempted the survey, and 89 completed the survey. List-wise deletion of participants was used to handle missing data when participants failed to answer more than 5 items in a row. To deal with randomly missing data, mean substitution was used. These GTAs were predominantly white (70%), domestic (73%), experienced (70%), PhD students (90%). Participants were evenly split between genders, with 55% identifying as female (see **Table 1)**.

**Table 1:** Summary of the demographics of Graduate Teaching Assistant (GTA) participants (n = 89 total), the calculated mean teaching anxiety with standard deviation, and the average self-efficacy (SE) scores and standard deviations across each subgroup. There were no significant differences in teaching anxiety among subgroups. Self-efficacy is measured on a 1-5 Likert scale, with 1 being “Not confident at all” and 5 being “Very confident”. Significant differences (p<0.05) are indicated with stars.

|  | n | % of total participants | Average teaching anxiety | SD for teaching anxiety | Average learning SE | SD for learning SE | Average instructional SE | SD for instructional SE |
| --- | --- | --- | --- | --- | --- | --- | --- | --- |
| **Gender** |  |  |  |  |  |  |  |  |
| Male GTAs | 40 | 45 | 65.8 | 15.7 | 3.9 | 0.55 | 3.9 | 0.68 |
| Female GTAs | 49 | 55 | 69.7 | 16.5 | 3.9 | 0.69 | 3.9 | 0.77 |
| **Citizenship Status** |  |  |  |  |  |  |  |  |
| Domestic | 65 | 73 | 69.0 | 15.6 | 3.8 | 0.61 | 3.8 | 0.71 |
| International | 24 | 27 | 65.1 | 17.6 | 3.9 | 0.66 | 4.1 | 0.79 |
| **Ethnicity** |  |  |  |  |  |  |  |  |
| White | 63 | 70 | 68.9 | 14.4 | 3.8 | 0.57 | 3.8 | 0.70 |
| Non-white | 26 | 30 | 65.5 | 19.7 | 3.9 | 0.76 | 4.1 | 0.80 |
| **Experience level** |  |  |  |  |  |  |  |  |
| Novice | 27 | 30 | 71.0 | 15.7 | 3.8 | 0.73 | 3.6* | 0.87 |
| Experienced | 62 | 70 | 66.6 | 16.5 | 3.9 | 0.58 | 4.0* | 0.64 |
| **Degree** |  |  |  |  |  |  |  |  |
| MS | 9 | 10 | 70.0 | 19.0 | 4.0 | 0.47 | 4.0 | 0.74 |
| PhD | 80 | 90 | 67.7 | 15.9 | 3.8 | 0.65 | 3.9 | 0.74 |
| **Department** |  |  |  |  |  |  |  |  |
| BCMB | 25 | 28 | 69.2 | 16.1 | 3.8 | 0.59 | 3.9 | 0.87 |
| EEB | 31 | 35 | 66.3 | 17.0 | 3.8 | 0.66 | 3.8 | 0.73 |
| GST | 9 | 10 | 58 | 15.1 | 4.2 | 0.55 | 4.3 | 0.64 |
| Micro | 23 | 26 | 72.6 | 14.4 | 3.8 | 0.60 | 3.9 | 0.62 |
| Other | 1 | 1 | 61 | N/A | 4.7 | N/A | 4.3 | N/A |

### Reliability of instruments

To assess the reliability of the hypothesized factors in our data, we calculated Cronbach alpha among our GTA population. We found for the teaching anxiety instrument, an alpha coefficient of 0.93; the self-efficacy instrument, an alpha coefficient of 0.88 for each construct; and lastly for the coping instrument alpha coefficients between 0.60-0.94 (*preparing materials* α = 0.60, *muscular desensitization* α = 0.78, *cognitive restructuring* α = 0.81, *preparing delivery* α = 0.88, *visualization* α = 0.94, and *mentoring* α = 0.80). Because the *preparing materials* (Coping 1) construct had poor reliability scores < 0.7, it was removed from further analysis.

### Teaching anxiety, self-efficacy, and coping among subgroups

Based on Parson’s (1973) *teaching anxiety* scale, an individual could score between 29 and 145. Average anxiety among the Biology GTAs was 67.4 (± 16.4 SD), indicating a mid-range level of anxiety. The minimum anxiety score for the sample was 32, and maximum was 116 (**Figure 1**). Average *self-efficacy* scores for both learning environment and instructional self-efficacy were 3.9 (± 0.64 SD), indicating higher than average perception of self-efficacy in teaching. Average coping frequency ranged from 3.62 to 23.24 depending on the coping strategy (see **Table 2** for GTA coping averages compared to their potential ranges).

**Table 2:**
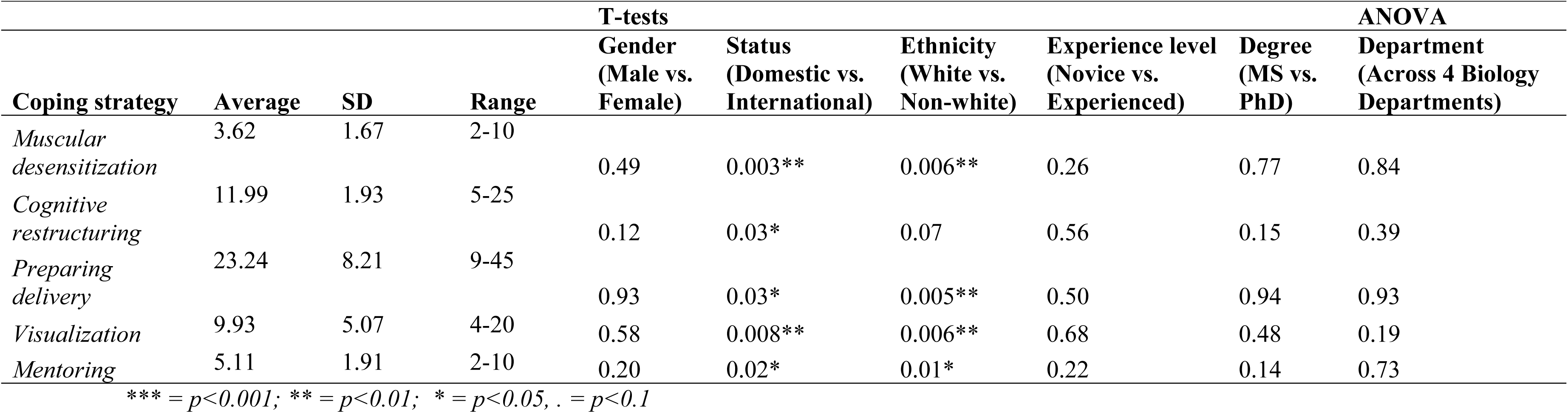
Calculated mean, standard deviation, and potential score range of coping strategies along with resulting p-values of t-tests and one-way ANOVAs among different subgroups within Biology GTAs participants (n=89). Significant differences were found among student Status and Ethnicity categories. Non-white, international students had significantly greater coping frequency than white, domestic GTAs. The five types of coping strategies for teaching anxiety kept in the analysis were: (1) *muscular desensitization*, (2) *cognitive restructuring*, (3) *preparing delivery*, (4) *visualization*, and (5) *mentoring*. Coping through *preparing materials* (Coping 1) was removed after the CFA due to low reliability.

**Figure 1:**
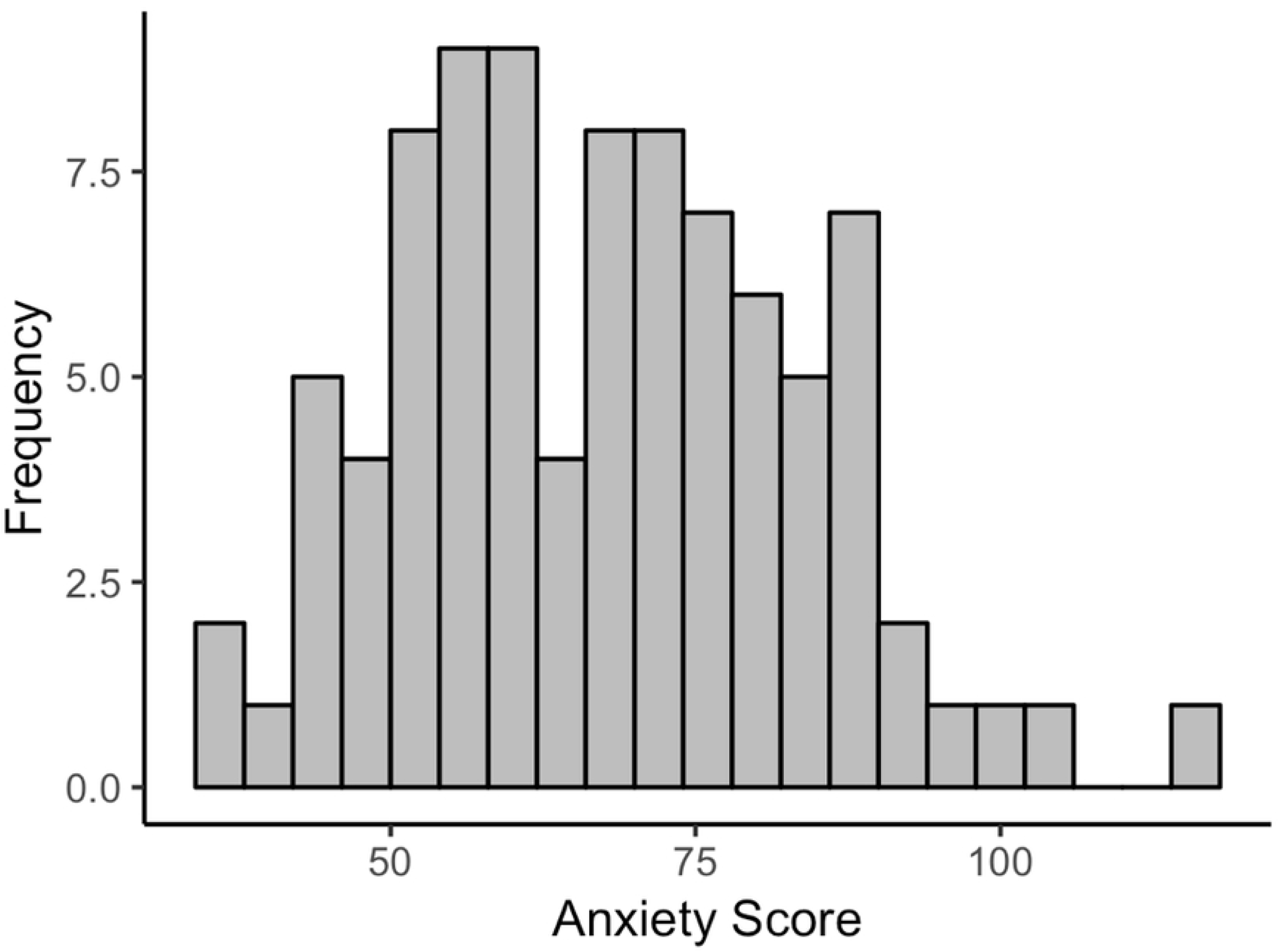
Distribution of teaching anxiety of GTA participants (n = 89). There is a relatively normal distribution of teaching anxiety in the GTA sample. There were 29 items in the anxiety measure, each rated on a 1-5 Likert scale. An individual could range between 29 to 145 on this anxiety scale, with 29 being the lowest level of anxiety and 145 being the highest.

#### Teaching Anxiety

There were no significant differences in teaching anxiety between GTAs of different genders, ethnicities, departments, year of study, or teaching experience level (**Table 1; Figure 2a**). However, examining the differences in general (not teaching) anxiety among subgroups, we found that international, non-white students had significantly less self-reported anxiety in graduate school generally (*t*=2.77, *p*<0.05) and in daily life (*t*=2.40, *p*<0.05).

**Figure 2:**
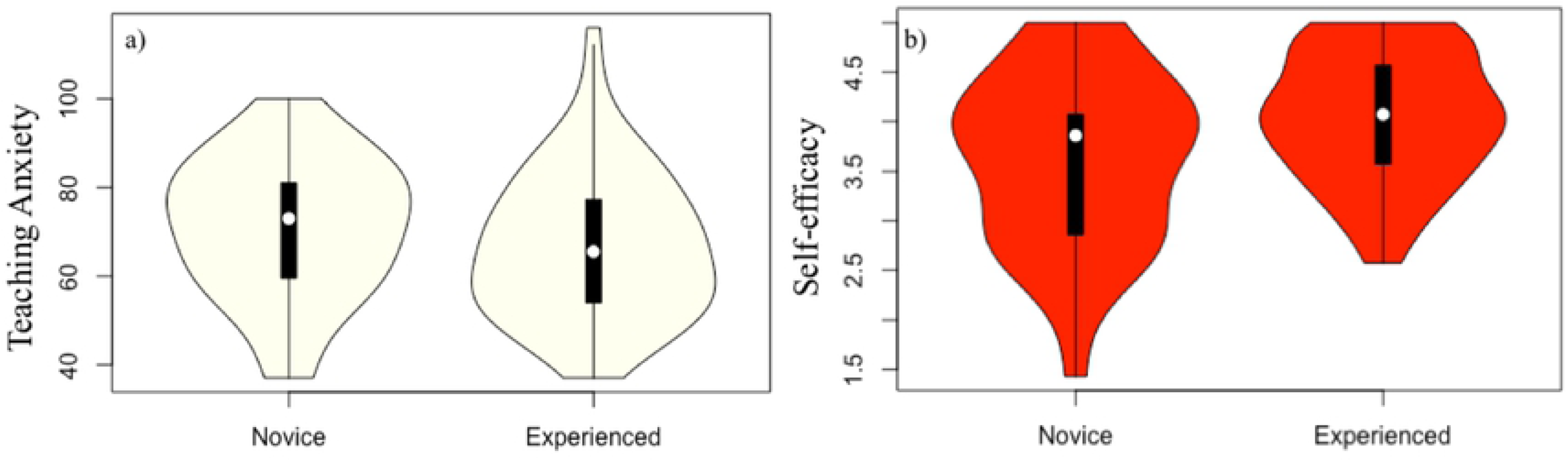
Differences in **a) teaching anxiety** (*t*=1.2, *ns*) and **b) instructional self-efficacy** (*t*=-2.3, *p*<0.05) between novice and experienced GTAs. Experienced GTAs (𝓍 =4.0±0.64) had significantly higher instructional self-efficacy than novice GTAs (𝓍 =3.6±0.87). There were no significant differences found between Novice and Experienced GTAs for the learning environment self-efficacy construct.

#### Self-efficacy

There were differences among the GTAs’ self-efficacy: experienced GTAs had significantly higher instructional self-efficacy (*t*=-2.28, *p*<0.05) than novice GTAs (**Table 1**; **Figure 2b**). This difference was not found in the learning environment self-efficacy construct. There were no other significant differences in teaching self-efficacy between other subgroups (**Table 1**).

#### Coping

Comparing differences in coping strategies between subgroups, we found significant differences between student citizen status subgroups (domestic vs. international) and Ethnic (white vs. non-white) groups (**Table 2**). There was high overlap between these two subgroups; 91% percent of the white GTA population were also domestic students, and 83% of the non-white GTAs were international students. Because of this overlap, we chose to compare only ethnicity to further examine trends in coping, since using both subgroup categories (Ethnicity and Status) would be highly redundant. We found significant differences between non-white (n=26) and white (n=63) GTAs (**Figure 3)**. Non-white groups coped significantly more often than their white counterparts. These coping strategies included **a) muscular desensitization** (*t*=2.93, *p*<0.001), **b) preparing delivery** (*t*=2.90, *p*<0.001), **c) visualization of oneself teaching successfully** (*t*=2.86, *p*<0.001) and **d) seeking mentoring (***t*=2.68, *p*<0.05; for further details about these coping strategies, see Methods).

**Figure 3:**
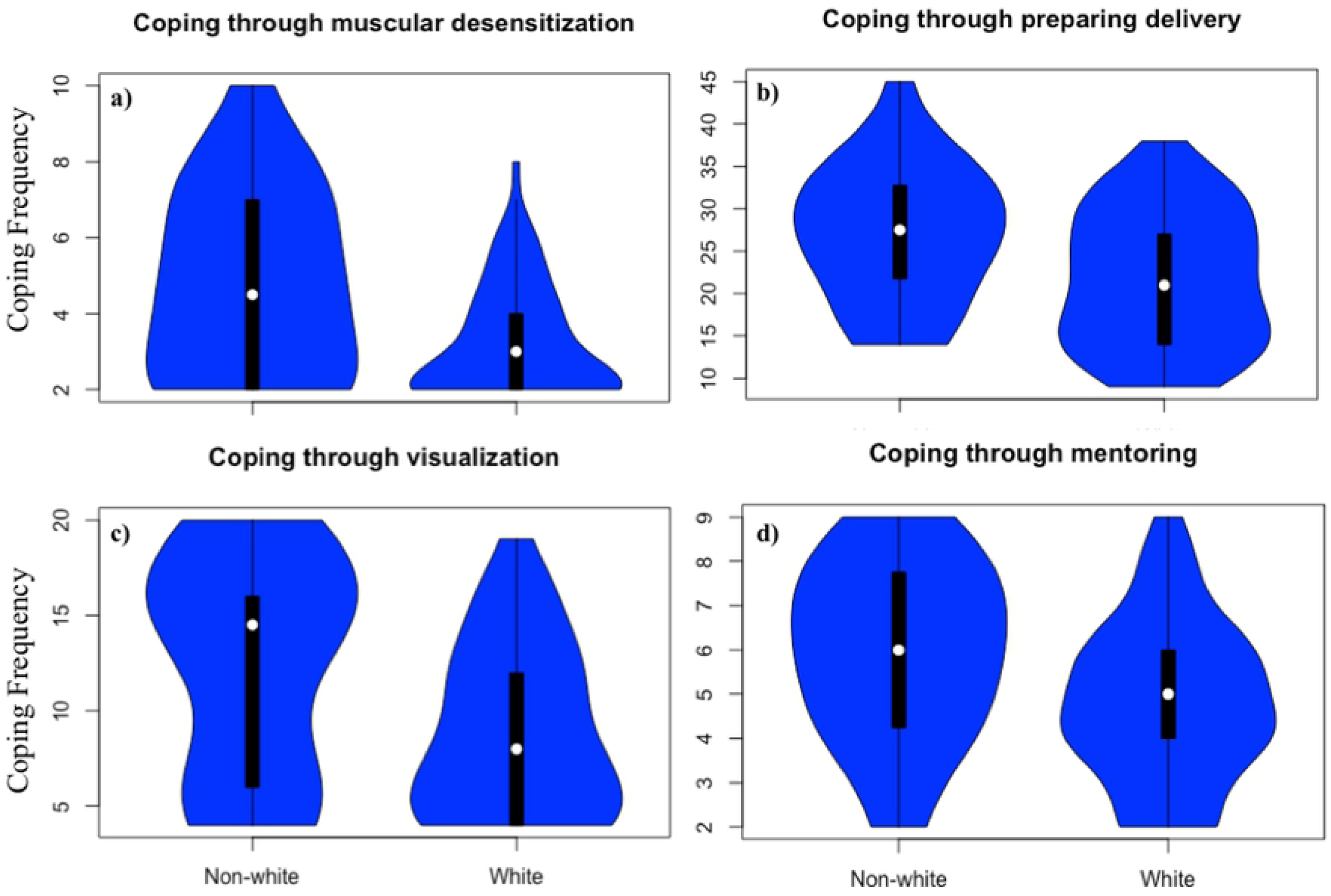
Differences in coping between non-white (n=26) and white (n=63) groups. Non-white groups coped significantly more than their white counterparts for these coping strategies. These coping strategies included **a) muscular desensitization** (Δ𝓍 =1.5, *t*=2.93, *p*<0.001), **b) preparing delivery** ((Δ𝓍 =5.4, *t*=2.90, *p*<0.0001), **c) visualization of oneself teaching successfully** (Δ𝓍 =3.5, *t*=2.86, *p*<0.001), **d)** mentoring (Δ𝓍 =1.2, *t*=2.68, *p*<0.05).

### Correlations of teaching anxiety, teaching self-efficacy, coping, and general anxiety

We next examined correlational relationships among variables as depicted in a correlogram (**Figure 4a and b**, R package “corrplot” [44]). These correlational results reveal associations between constructs, which provide a preliminary indication of the statistical significance, strength, and the direction (positive or negative) of these associations. We found that both constructs of teaching self-efficacy were significantly and negatively associated with teaching anxiety (**Figure 4a**, *r = −0.59, p<0.05*). Coping strategies had significant strong to moderate correlations among other coping strategies (**Figure 4a**, *r = 0.30-0.60, p<0.05*) and moderate to weak correlations with self-efficacy constructs (**Figure 4a**, *r = 0.04-0.40, p<0.05*).

**Figure 4:**
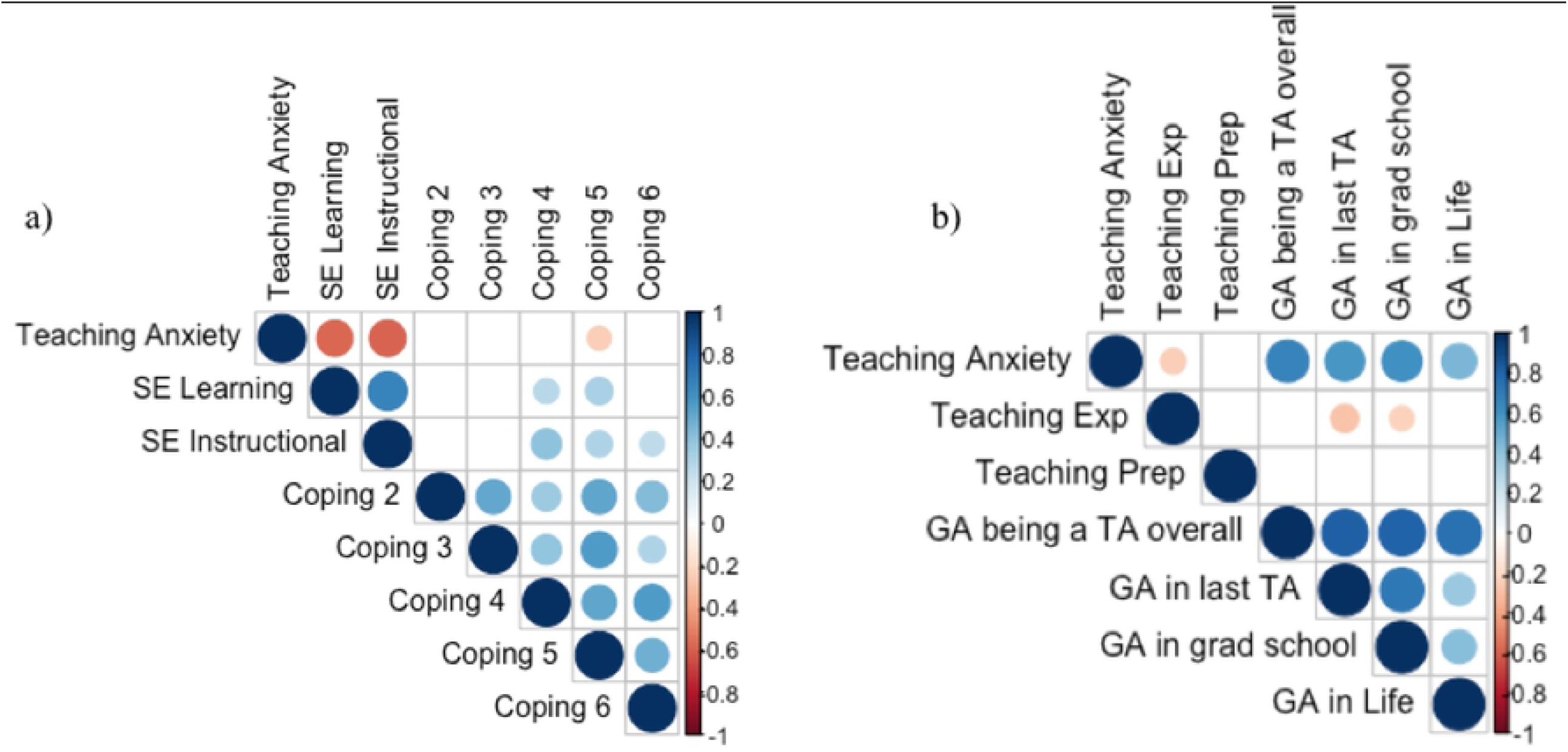
Correlograms of bivariate correlations among **a)** study constructs: teaching anxiety, teaching self-efficacy, and coping strategies (N=89). Coping 1 to 6 strategies are as follows: *preparing materials, muscular desensitization, cognitive restructuring, preparing delivery, visualization*, and *mentoring*. Coping 1 was taken out because of poor reliability scores. The second correlogram depicts correlations between **b)** teaching anxiety and contextual variables (total semesters of teaching experience, total hours of teaching preparation, and four general anxiety (GA) items related to general anxiety about being a graduate student, being a TA in a GTAs most recent teaching assignment, being a GTA generally, and general anxiety in their daily life). Positive correlations are displayed in blue and negative correlations in red color. Correlation coefficients are proportional to the color intensity and the size of the circle. The legend color shows the correlation coefficients according to the corresponding colors. Correlations with p-value > 0.05 are considered as insignificant and have a blank and no circle.

In correlations between teaching anxiety and continuous background variables (total semesters of teaching experience, hours of teaching preparation, and general anxiety items), we found that general anxiety had a significant positive relationship with teaching anxiety (**Figure 4b**, *r = 0.46-0.67, p<0.05*). Total semesters of teaching experience were also weakly negatively correlated to teaching anxiety (**Figure 4b**, *r = −0.24, p<0.05*), general anxiety in a GTA’s last teaching assignment (**Figure 4b**, *r = −0.29, p<0.05*), and general anxiety in graduate school (**Figure 4b**, *r = −0.22, p<0.05*).

### Model for teaching anxiety

Using multiple linear regressions, we found that both learning and instructional self-efficacy significantly predicted GTA teaching anxiety, explaining 66% of the variance, along with two items measuring general anxiety: general anxiety as a GTA, and anxiety related to their last teaching assignment (**Table 3a**, *R*^*2*^_*adi*_ = 0.66, *p* <0.001, AIC=164). A second model was developed removing general anxiety measures, as that construct was measured using four non-validated investigator-created questions (**Table 3b**, *R*^*2*^_*adi*_ = 0.46, *p* <0.001, AIC=208). This second model did not capture as much variance as the first. For the second model, the step-wise selection procedure chose 9 variables, 7 of which were significant in explaining the variance in teaching anxiety. These variables included both teaching self-efficacy constructs, teaching experience, and 4 types of coping measures (coping through preparing delivery, cognitive restructuring, and mentoring). The non-significant variables included hours of teaching preparation and feelings of departmental support. This model explained 46% of variance in teaching anxiety. Both models of teaching anxiety had no significant multicollinearity within the model. Comparing models using the AIC scores, we found that the initial model with general anxiety included was more parsimonious. VIF calculations revealed no inflation issues due to multicollinearity.

**Table 3:**
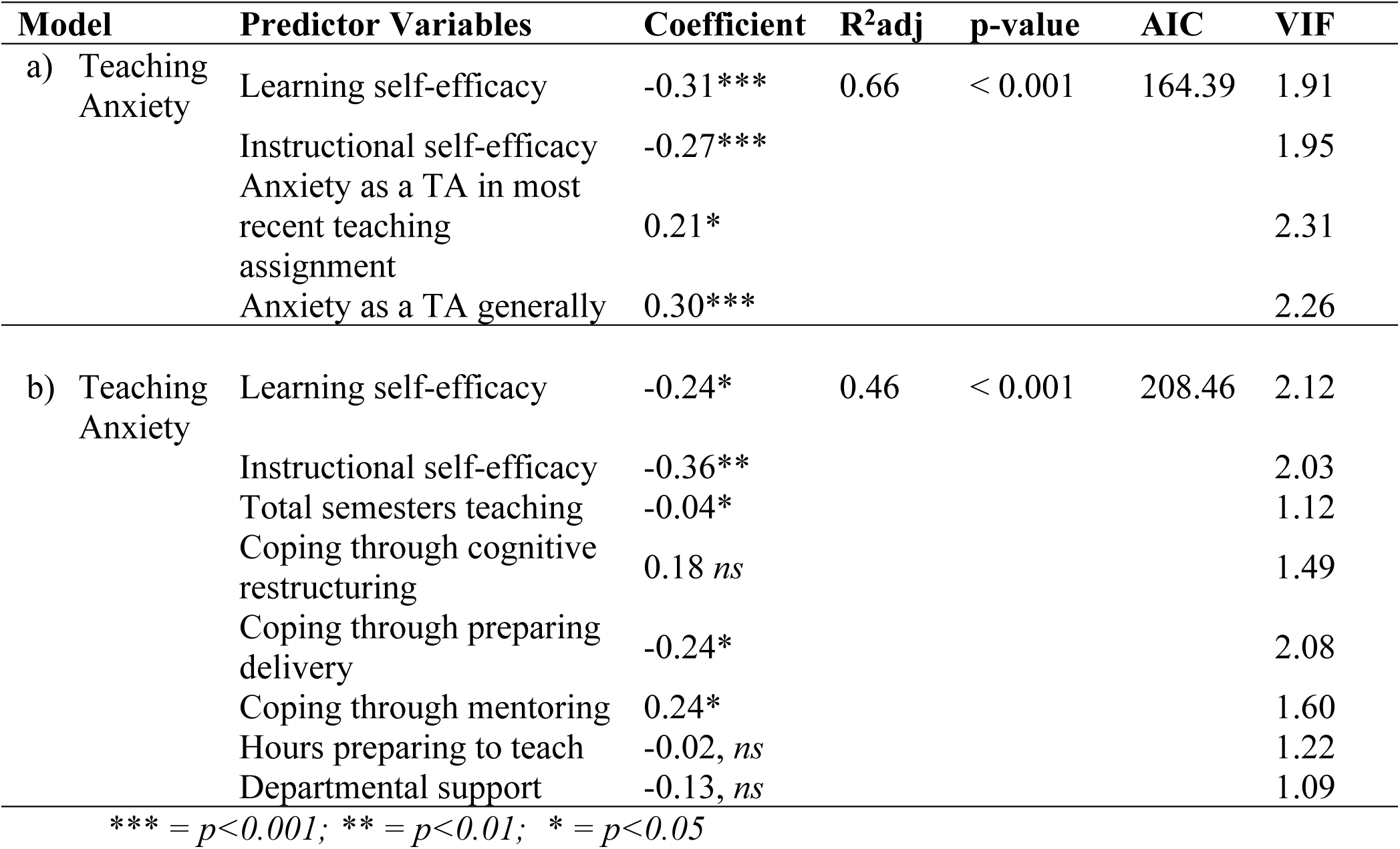
Multiple linear regressions were built to a) determine what variables contributed to GTA teaching anxiety in Fall 2016 including general anxiety and b) without general anxiety. Both models were significant, explaining over 46% of the variance in teaching anxiety (n=89). The first model is the most parsimonious, with 66% of the variance in teaching anxiety explained. Instruments were z-scored for teaching anxiety, self-efficacy, and coping scores, to facilitate comparisons across instruments.

## DISCUSSION

In answering the research questions related to this study, we found that teaching self-efficacy plays an integral role in reducing teaching anxiety in Biology GTAs at our institution, and that coping frequency may contribute to building teaching self-efficacy. Perhaps unsurprisingly, GTA general anxiety was positively related to teaching anxiety, although those results should be treated with caution because of the unvalidated nature of the general anxiety items. Teaching anxiety appeared as a normal distribution in our population, with the majority of GTAs reporting moderate levels of anxiety. This anxiety is universal among subgroups, with similar levels of anxiety across genders, ethnicities, student status, and experience level. Interestingly, what significantly differed among subgroups was teaching self-efficacy and frequencies of different coping strategies. Experienced GTAs had significantly greater instructional self-efficacy than novice GTAs. Non-whites and international GTAs had greater coping frequency than their white, domestic student counterparts. Since we did not find a significant difference in teaching self-efficacy among those groups, effective coping strategies may be contributing to the lack of anxiety differences.

### Teaching anxiety levels were mostly similar among GTAs, but general anxiety differed

Despite evidence suggesting female graduate students suffer higher rates of general anxiety and depression than male graduate students [2], we did not find gender differences in teaching anxiety. Teaching is a role often dominated by women, especially in primary and secondary education [45]. Women gravitate toward teaching-centered occupations more often than men, with sometimes greater self-efficacy for the task compared to their male counterparts [46,47]. When comparing how gender role socialization might contribute to gender differences in self-efficacy and confidence, Betz and Hackett [46] found that women demonstrated significantly greater self-efficacy for traditionally female occupations and much lower efficacy for traditionally male occupations compared to men. These trends in self-efficacy between genders, however, have not always been consistently observed [24,48,49] More recently, when comparing effects of teaching professional development (PD) between genders, [24] found female graduate students had lower self-efficacy than male GTAs when both groups lacked any PD experience. Interestingly, this gap became significantly smaller as women became more engaged in teaching PD activities. In our study population, GTAs are supported by many teaching PD opportunities at the institutional and departmental level, possibly increasing self-efficacy and decreasing teaching anxiety in female GTAs [25]. Though we did not explicitly ask about the intensity of their PD participation, 70% of the study participants were *experienced* GTAs, making the likelihood of GTAs having participated in PD (via CIRTL programs, early-semester orientation or course preparation meeting, and workshops let by institutional Teaching and Learning programs) higher.

Though no teaching anxiety differences were found among other subgroups, international, non-white students were found to have significantly less general anxiety than their domestic, white student counterparts. We had predicted that those not acclimated to Western cultures and languages would be more anxious teaching [34,50–52], however, our data suggested no discernable differences between these groups. International, non-white GTAs had significantly greater frequency of coping strategies than domestic, white GTAs. Therefore, a lack of teaching anxiety difference between ethnic and student status groups may be attributed to more effective coping strategies employed by international, non-white GTAs. When Roach and Olaniran [51] studied 201 international graduate students across multiple disciplines, they also found that international GTAs had low levels of intercultural communication apprehension or anxiety, expressing a great willingness to communicate with people from a different culture.

### Teaching self-efficacy varies with experience level

Unsurprisingly, experienced GTAs had significantly higher instructional self-efficacy compared to novice GTAs. As indicated in the introduction, self-efficacy may be built by four main mechanisms: mastery experiences, vicarious experiences, social persuasion, and emotional or physiological arousal. Bandura [53] purported that mastery experience was the strongest and most influential in building self-efficacy in a task. Mastery experiences require individuals to directly encounter and conduct a given task, therefore GTAs who have greater authentic classroom teaching experiences (e.g. guest lecture, instructor of record, not just a grader), would have higher teaching self-efficacy. In our study, this difference in self-efficacy between novice and experienced GTAs was only found for the *instructional self-efficacy* construct. Instructional self-efficacy is related to activities needed to prepare and teach a class, while learning environment self-efficacy focuses on tasks related to promoting and providing a positive, engaging, and respectful classroom environment [11,12]. Morris and Usher [54] studied the teaching self-efficacy of award-winning professors, and determined that early successful instructional experiences, which are a combination of mastery experiences and verbal persuasions, are important for developing high teaching self-efficacy. Interestingly, our data reveal which *type* of teaching self-efficacy Biology GTAs appear to build with greater experience initially: instructional self-efficacy.

### Building greater teaching self-efficacy may reduce teaching anxiety

Both self-efficacy constructs and general anxiety significantly contributed to teaching anxiety, with self-efficacy constructs negatively correlated to teaching anxiety i.e. the higher teaching anxiety, the lower the reported teaching self-efficacy. This relationship between anxiety and self-efficacy broadly has been well-established in social cognitive theory. According to Bandura [4,53] possessing self-efficacy or a perceived self-efficacy to control potential external stressors or threats, plays a central role in anxiety arousal. Individuals who believe they can exercise control over these potential threats do not have apprehensive cognitions and thus, are not disturbed by them. However, individuals who cannot manage or perceive they cannot manage potential stressors experience high levels of anxiety arousal. Bandura [4] states that this latter group tends to dwell on their coping deficiencies and view many aspects of their environment as holding potential anxiety-provoking danger. Thus, perceived control or self-efficacy over a stressor, even if not substantiated in actuality, reduces anxiety. The question that naturally emerges from such a result is what then contributes to building self-efficacy? What Bandura (1988) suggests is that coping is a type of self-efficacy necessary in moderating anxiety.

Previous structural equation models have been developed for teaching self-efficacy constructs for STEM GTAs [12]. Those models revealed the importance of K–12 teaching experience, hours and perceived quality of GTA PD, and perception of support by the department in developing teaching self-efficacy. These models, however, did not consider how teaching anxiety and alleviating teaching anxiety through coping may also contribute to self-efficacy. As the growing evidence suggests [24,25,32,55,56], teaching PD opportunities allow GTAs to build self-efficacy and reduce anxiety in teaching. Reeves et al. [25] examined the impact of GTA training programs at three separate institutions and determined that regardless of the differences in program settings, PD was associated with gains in content knowledge and self-efficacy, and decreases in teaching anxiety. By advocating for quality PD opportunities for GTAs, coping efficacy for teaching anxiety may be strengthened.

### Coping with teaching anxiety may contribute to building teaching self-efficacy

Effective coping of a potential stressor or threat must precede any increase in self-efficacy for a task, especially if there is anxiety towards said task. According to Skinner et al. [28], there are 12 families of coping, which can be further categorized as *adaptive* or *maladaptive* [30]. Adaptive coping helps individuals successfully progress through problems and supports their well-being; while maladaptive coping prevents stressors or problems from being solved and can threaten well-being. When examining which of the families these preparation coping strategies (*i.e.* preparing materials and preparing delivery) align with most, *problem-solving* and *information-seeking* both require planning or preparing as a response to external stressors. Problem-solving attempts to resolve the stressor, through planning and/or enacting a potential solution, while information-seeking attempts to learn more about the stressor. When comparing coping among subgroups, along with preparation coping strategies, international, non-white GTAs also had significantly greater visualization of oneself successfully teaching. This strategy could be categorized under Skinner et al.’s *self-reliance* family, where cognitive or emotional regulation is used to perceive the stressor more positively. Coping through preparing materials, coping through preparing delivery, and visualization of oneself successfully teaching, all fall under *adaptive* coping [28,30]. These types of approach-orientated, adaptive coping can lead to positive increases of self-efficacy and reductions in anxiety.

### Implications for GTA professional identity, attrition, and graduate student well-being

Teaching anxiety is one facet contributing to overall graduate student well-being. Generally, feelings of anxiety can escalate into diagnosed anxiety disorder, which impedes functioning in daily life. The anxieties graduate students face affect not only their overall mental health, but also reduce their retention in graduate programs and academia [1,57–59]. Some of this anxiety may be attributed to the multiple responsibilities of graduate students. Graduate students are professionals-in-training, balancing simultaneous roles as teachers, researchers, students, and employees [60,61].

During graduate school, students are in a state of transition where they experience a variety of identities and roles [60–62]; anxiety over role responsibilities can be expected. Though we have only addressed anxiety related to teaching, research anxiety or anxiety related to conflicting roles as a graduate student has yet to be studied. Some would argue that research is the primary identity in which graduate students must develop during graduate school; with teaching being perceived, at best, as a resume builder, and at worst, as a punishment for those unable to acquire fellowships for teaching releases [63,64]. Recent data, however, suggest that graduate student investment of time in PD may be beneficial for research preparation [65]. Examining how research anxiety relates to teaching anxiety, how these anxieties change over time (especially as GTAs grow in mastery experiences), and *why* GTAs are anxious needs further exploration.

The level of anxiety experienced by individual graduate students may contribute to who persists in academic careers and how they perform in their jobs if they do persist. Non-tenure track faculty, for example, also report anxiety, depression [66] and burnout [67,68]. This anxiety can also negatively impact student learning [14,69]. Research on K-12 classrooms has found negative relationships between teacher anxiety and grading practices [69], rapport and interpersonal relationships with students [70], and academic performance [71]. Equipping our future Biology faculty with the tools to discuss anxiety and ways to cope, may improve the success of future instructional faculty.

### Limitations

As with all studies, the results must be interpreted in light of our limitations. The results of this study are not generalizable, as we sampled from a self-selected pool of Biology GTAs from one institution. Three main methodological limitations also restrict our ability to generalize results to a wider population, including: 1) self-reporting of anxiety, 2) using investigator-created items for general anxiety, and 3) measuring frequency of coping.

Critics of measuring teaching anxiety through self-reporting assert that there is poor evidence to suggest there is any influence in teaching performance [72] or it conflates teaching anxiety with merely “teaching concerns” [73]. However, teaching anxiety may be perceived to some extent by an external observer [36,55,69]. To test this, Parson [36] had 25 preservice teachers score their own teaching anxiety and then correlated those scores to ratings provided from teacher supervisors’ after a teaching observation. They found evidence suggesting that the teaching anxiety reported on the scale corresponded to what may be externally perceived teaching anxiety (r = 0.24-0.54). Though Parsons’ instrument was developed to measure teaching anxiety in preservice K-12 teachers, her instrument has been implemented widely in many other K-12 and college teacher populations with similar distributions of teaching anxiety across the scale [32,55,69].

The second limitation pertains to the items in which we measured general anxiety among GTAs. As we mentioned in the Methods, because our sample size did not allow for any of our measures to undergo confirmatory factor analysis [37], we were unable to even initially test whether the general anxiety measures we used formed a true “general anxiety” factor. Thus, although they were useful in exploring potential relationships between different types of anxiety, these results should be treated with caution. Further evidence must be collected to support general anxiety differences among ethnic and student status groups or relationships with teaching anxiety.

The third limitation involves measuring coping. Bandura [4] indicated coping efficacy was an integral component in exercising control over anxiety arousal. However, in this study, the instrument we used measured coping frequency [14]. Though coping frequency can be used as a marker for coping strategies, it is not equivalent to measuring efficacy of coping. Roach [14] agrees that it is possible that TAs with high teaching anxiety could spend hours preparing for a class and still be unsuccessful because of *how* they prepared or the efficacy of coping. Roach [14] suggests that this is where the GTA supervisor or teaching mentor must help the GTA make more efficacious coping decisions. Future work could attempt to capture the effectiveness of coping strategies enacted instead of only frequencies.

### Conclusions

To tackle the anxiety epidemic in academia, particularly in regards to graduate student teaching, there must be opportunities for self-efficacy to be built and anxiety to be reduced. Teaching PD activities or training opportunities for GTAs pose a tangible, effective method for institutions and departments to consider to help reduce anxiety and increase self-efficacy. PD workshops may also provide efficacious coping strategies to regulate external stressors which cause anxiety, especially to particular graduate student subgroups. We suggest further research to explore whether lack of teaching anxiety differences among gender and ethnic subgroups may be attributed to PD. Projects focused on PD of graduate students, such as the Biology Teaching Assistant Program (BioTAP) or the Longitudinal Study of Future STEM Scholars (LSFSS), can further the scholarship necessary to understand the relationships among graduate students, teaching, anxiety, and coping.

## ACKNOWLEDGEMENTS

Thank you to Caroline Wienhold, Benjamin England, Margaurete Romero, Sarah Andrews, Nicole Chodkowski, Robert Furrow, Nicholas Nagle, Kimberly Sheldon, Randall Small, and Erin Hardin for their comments on earlier versions of the manuscript. Thank you to all GTA participants for taking part in this research.

## SUPPORTING INFORMATION

**S1 Table. Biology GTA (n=89) survey data.** All identifying information has been removed. List-wise deletion of participants was used to handle missing data when participants failed to answer more than 5 items in a row. Mean substitution was used to handle randomly missing data.

**S2 Table: Survey provided to Biology GTAs in Fall 2016.** Corresponding codes to the data for answers to each question are presented in parentheticals.

